# A BONCAT-iTRAQ method enables temporally resolved quantitative profiling of newly synthesised proteins in *Leishmania mexicana* parasites during starvation

**DOI:** 10.1101/713735

**Authors:** Karunakaran Kalesh, Paul W. Denny

**Affiliations:** Department of Chemistry, Durham University, Durham, United Kingdom; Department of Biosciences, Durham University, Durham, United Kingdom

## Abstract

Adaptation to starvation is integral to the *Leishmania* life cycle. The parasite can survive prolonged periods of nutrient deprivation both *in vitro* and *in vivo* and starvation plays a crucial role in the differentiation of the parasite from the non-infective promastigote form to infective metacyclics. The identification of parasite proteins synthesised during starvation is key to unravelling the molecular mechanisms facilitating adaptation to these conditions and the associated lifecycle differentiation. Additionally, as stress adaptation mechanisms in *Leishmania* are linked to virulence as well as infectivity, profiling of the complete repertoire of Newly Synthesized Proteins (NSPs) under starvation is important for drug target discovery. However, differential identification and quantitation of low abundance, starvation-specific NSPs from the larger background of the pre-existing parasite proteome has proven difficult, as this demands a highly selective and sensitive methodology. Herein we introduce an integrated chemical proteomics method in *L. mexicana* promastigotes that involves a powerful combination of the BONCAT technique and iTRAQ 4-plex quantitative proteomics Mass Spectrometry (MS), which enabled temporally resolved quantitative profiling of *de novo* protein synthesis in the starving parasite. Uniquely, this approach integrates the high specificity of the BONCAT technique for the NSPs, with the high sensitivity and multiplexed quantitation capability of the iTRAQ proteomics MS. Proof-of-concept experiments identified a total of 166 NSPs in the parasite and quantified the relative changes in abundance of these proteins as a function of duration of starvation. Our results show a starvation time-dependent differential expression of important translation regulators. GO analysis of the identified NSPs for Biological Process revealed translation (enrichment P value 6.93e^−67^) and peptide biosynthetic process (enrichment P value 1.85e^−66^) as extremely significantly enriched terms indicating the high specificity of the NSP towards regulation of protein synthesis. We believe that this approach will find wide-spread use in the study of the developmental stages of *Leishmania* species and in the broader field of protozoan biology.

**Author Summary:** Periodic nutrient scarcity plays crucial roles in the life cycle of the protozoan parasite *Leishmania spp*. Starvation triggers differentiation of the parasite from a non-infective form to an infective form. Although adaptation to nutrient stress has a pivotal role in *Leishmania* biology, the underlying mechanisms remain poorly understood. In a period of nutrient starvation, the parasite responds by decreasing its protein production to conserve nutrient resources and to prevent formation of toxic proteins. However, even during severe starvation, the parasite generates certain essential quality control and rescue proteins. Differential identification of the complete repertoire of these proteins synthesised during starvation from the pre-existing proteins in the parasite holds the key to understanding the starvation adaptation mechanisms. This has been challenging to accomplish due to technical limitations. Using a combination of chemical labelling techniques and protein mass-spectrometry, we selectively identified and measured the proteins generated in the starving *Leishmania* parasite. Our results show a starvation time-dependent differential expression of important protein synthesis regulators in the parasite. This will serve as an important dataset for a holistic understanding of the starvation adaptation mechanisms in *Leishmania*. We also believe that this method will find wide-spread applications in the field of protozoa and other parasites causing Neglected Tropical Diseases.

## Introduction

Protozoan parasites of the *Leishmania spp.* are the causative agents of leishmaniasis, a Neglected Tropical Disease (NTD) endemic in over 90 countries worldwide, affecting approximately 12 million people with an estimated 700,000 to 1 million new cases annually [1]. These protozoa have a complex life cycle, progressing from extracellular promastigote stages in the sandfly vector to the obligate intramacrophage amastigote stage in the mammalian host [2]. During their digenetic life cycle *Leishmania spp.* are exposed to extreme stress conditions, including severe nutrient starvation, and the parasites have evolved mechanisms to adapt to and surmount large fluctuations in nutrient availability [3–5]. Nutrient starvation is also known to be critical for metacyclogenesis, a process that involves differentiation of non-infective procyclic promastigotes to the infective metacyclic promastigote stage in the sandfly vector [6,7]. However, the key proteins involved in the starvation-adaptation mechanisms of the parasite remains unknown and the identification and quantitation of proteins synthesised *de novo* during starvation is critical to develop understanding of these. Progress in this direction has been hampered by technical limitations; the lack of availability of a robust and sensitive method that can differentiate and characterise the *de novo* synthesised proteins during starvation from the complex cellular background of pre-existing proteome being the major bottleneck. Herein we describe a combination of the bio-orthogonal non-canonical amino acid tagging (BONCAT) [8,9] technology and isobaric tags for relative and absolute quantification (iTRAQ) quantitative mass-spectrometry (MS) proteomics [10,11] to quantitatively profile the newly synthesised proteome (NSP) of *Leishmania mexicana* promastigotes during starvation.

Regulation of eukaryotic gene expression involves a coordinated network of molecular processes staring with initiation of transcription, followed by post-transcriptional regulatory mechanisms. Processing of the primary transcript-RNAs essentially involves three steps namely capping, where a 7-methylguanosine moiety modifies the 5’ end, and polyadenylation, where a poly-A tail is added at the 3’ end, and finally removal of the intron sequences via splicing. The processed RNAs (mRNAs) are then transported to the cytoplasm for the translation to take place. The transcripts interact with several proteins to form messenger ribonucleoprotein complexes (mRNPs), which regulate many aspects of mRNA stability and function. Intriguingly, regulation of gene expression in *Leishmania spp.*, and other similar kinetoplastids, is fundamentally different from other eukaryotes [12–14]. As the open reading frames of genes in these parasites are arranged in long polycistronic clusters, RNA polymerase II-dependent regulation of transcription initiation does not occur and instead, monocistronic mRNAs are generated by a trans-splicing mechanism and polyadenylation. The gene expression in *Leishmania spp.* is regulated almost exclusively at the post-transcriptional level and this involves protein-mediated molecular mechanisms controlling the mRNA degradation, RNA editing in the kinetoplast and protein translation. Because of the existence of an independent layer of translational regulation, the mRNA levels in *Leishmania* are not a good predictor of protein expression, and this poor correlation between the transcript and protein expressions warrants in-depth protein-level studies in this organism [15,16].

The proteome of an organism is highly dynamic, and the protein turnover is tightly controlled by multiple check points in protein synthesis and degradation processes. This dynamic protein turnover plays a crucial role in maintaining the cellular homeostasis. Cells respond to stimuli and perturbations by altering their protein expression levels. Measuring these changes in the proteome is important to understanding the underlying biological processes. MS proteomics serves as a powerful technique for directly measuring the effect of perturbations on cellular proteome [17]. However, during starvation in *Leishmania*, the global protein synthesis operates at a lower rate as the parasite tries to conserve available limited nutrient resources, a highly robust and sensitive enrichment method has to be coupled with the MS in order to differentially identify the NSPs from the larger background of the parasite’s existing proteome.

We envisaged that the BONCAT technique could be applied for a selective profiling of NSPs in *L. mexicana* parasites during starvation. BONCAT involves metabolic incorporation of a methionine surrogate non-canonical amino acid bearing a small bio-orthogonal functional group, such as L-azidohomoalanine (AHA) or L-homopropargylglycine (HPG) (Fig 1A), into the newly synthesised proteins (NSPs). As AHA and/or HPG are methionine surrogates, they are readily and efficiently incorporated into NSPs by cell’s endogenous translational machinery [8]. In this case, the presence of the bio-orthogonal click tag in the newly translated proteins provide an efficient means to distinguish, and selectively isolate these proteins from the pre-existing pool of proteins via a highly efficient copper (I) catalysed azide-alkyne cycloaddition (CuAAC) click reaction [18] with a capture reagent bearing the orthogonal functionality to the tag (alkyne vs. azide and *vice versa*). Additionally, in order to get a temporally resolved quantitative information of the effect of starvation on the protein synthesis in the parasite, we decided to couple the BONCAT approach with iTRAQ quantitative proteomics MS [10]. The iTRAQ, similar to the tandem mass tag (TMT) quantitative proteomics [19], is a peptide-level labelling approach that offers sample multiplexing. Importantly, the multiplexed isobaric tags provide an advantage of pooling of peptide signals across the different test conditions, which increases the sensitivity of detection of even low-abundant peptides [20,21]. Using a combination of BONCAT and iTRAQ-4plex quantitative MS, we have identified and quantified a total of 166 NSPs in *L. mexicana* promastigotes under starvation. The iTRAQ 4-plex platform enabled profiling of relative quantitative changes in the abundance of the NSPs at three different time points after the onset of starvation in the parasite. Subsequent gene ontology (GO) analysis of the data sets revealed significant enrichment of proteins involved in regulating protein translation in the parasite. This is the first quantitative proteomics study that revealed the NSPs along with their temporally resolved quantitative expression changes during starvation in *Leishmania*.

**Fig 1.**
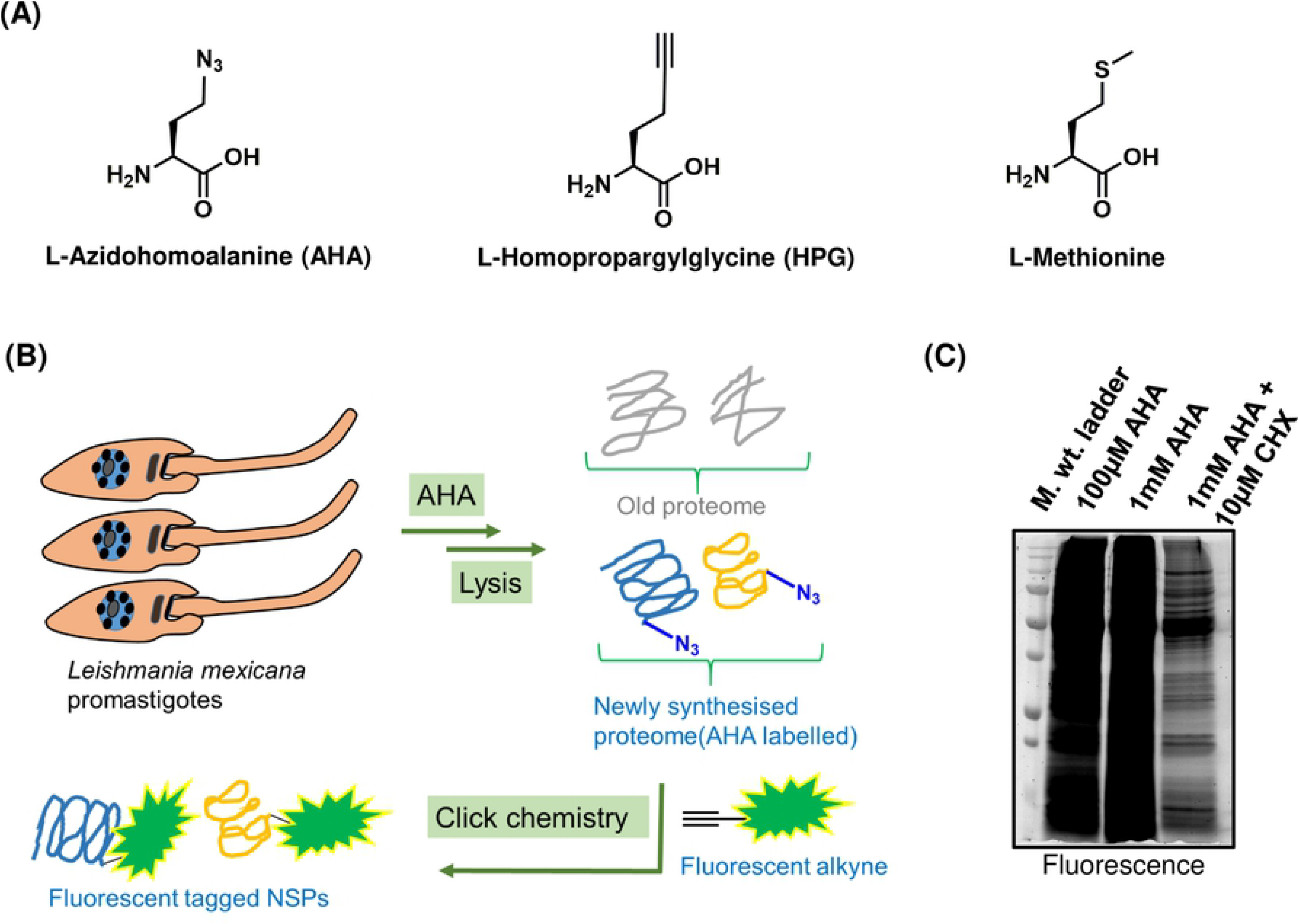
BONCAT in *L. mexicana* promastigotes. (**A**) Chemical structure of AHA, HPG and Methionine. (**B**) Workflow for BONCAT in *L. mexicana* promastigotes. AHA that can be bioorthogonally tagged with a fluorescent terminal alkyne was used for the BONCAT. (**C**) Fluorescent labelling of the NSPs following BONCAT detected by in-gel fluorescence scanning.

## Material and methods

### Chemicals and reagents

L-Azidohomoalanine (AHA; Iris Biotech GmbH), Cycloheximide (CHX; ACROS Organics), Tris (2-carboxyethy)phosphine hydrochloride (TCEP; Sigma Aldrich), 5-Tetramethylrhodamine-Alkyne (TAMRA-Alkyne; Jena Bioscience), Acetylene-PEG4-Biotin (Biotin-Alkyne; Jena Bioscience), Tris [(1-benzyl-1H-1,2,3-triazol-4-yl)methyl]amine (TBTA; Sigma Aldrich), Dimethyl sulfoxide (DMSO; Sigma Aldrich), Copper sulphate (CuSO_4_; Sigma Aldrich), 2-Amino-2-(hydroxymethyl)-1,3-propanediol (Tris; Sigma Aldrich), 4-(2-Hydroxyethyl)piperazine-1-ethanesulfonic acid (HEPES; Sigma Aldrich), Sodium chloride (NaCl; Fisher Scientific), Sodium dodecyl sulphate (SDS; Fisher Scientific), Sodium bicarbonate (NaHCO_3_; ACROS Organics), Calcium chloride (CaCl_2_; Sigma Aldrich), Urea (Sigma Aldrich), 1,4-Dithiothreitol (DTT; Sigma Aldrich), 2-Chloroacetamide (CAA; Sigma Aldrich), L-Glutamine solution (Sigma Aldrich), Benzonase (Sigma Aldrich), DC™ Protein Assay (Bio-Rad), Dialysed Foetal Bovine Serum (FBS; Life Technologies), Schneider’s Insect Medium (Sigma-Aldrich), Schneider’s Drosophila Medium without L-Methionine (PAN Biotech), Dulbecco Phosphate Buffered Saline (DPBS, Gibco), 1M Triethylammonium bicarbonate (TEAB) buffer (Sigma Aldrich), NeutrAvidin-Agarose beads (Thermo Scientific), iTRAQ® Reagents Multiplex Kit (Sigma Aldrich), Optima™ LC-MS Grade Trifluoroacetic acid (TFA; Fisher Scientific), Optima™ LC-MS Grade Formic acid (TFA; Fisher Scientific), Optima™ LC-MS Grade Acetonitrile (CAN; Fisher Scientific), Optima™ LC-MS Grade Methanol (MeOH; Fisher Scientific) and Sequencing Grade Modified Trypsin (Promega).

### Culturing of *L. mexicana* promastigotes

Promastigote form of *L. mexicana* strain M379 (MNYC/BC/62/M379) were grown in T-25 flasks at 26 °C in Schneider’s Insect Medium (Sigma-Aldrich) supplemented with 0.4g/L NaHCO_3_, 0.6g/L anhydrous CaCl_2_ and 10% FBS (pH 7.2).

### Metabolic labelling of newly synthesised proteins in *L. mexicana* promastigotes

The promastigotes in T-25 flasks were grown to mid log phase (~5 × 10^6^ parasites/mL) and washed with methionine-free Schneider’s medium supplemented with 10% dialysed FBS. The parasites were then incubated with methionine-free Schneider’s medium supplemented with 10% dialysed FBS for 30 minutes to deplete the intracellular methionine reserves. The parasites were labelled with AHA (100µM and 1mM) in fresh methionine-free Schneider’s medium supplemented with 10% dialysed FBS for 1 hour with or without CHX (10µM). In order to induce starvation, the parasites, after the initial 30 minutes of methionine depletion, were incubated with DPBS for different duration (1 hour to 7 hour) and treated with AHA (50µM) in DPBS for 1 more hour. DMSO was used as a vehicle control for the AHA treatment. In order to probe the NSPs since the point of the onset of severe starvation, in one of the samples the 1 hour AHA incubation was carried out concurrently with the first 1 hour DPBS treatment. This condition is defined as the 1 hour starvation in the experiments. Following the AHA treatment, the parasites were lysed using lysis buffer (50mM HEPES, pH 7.4, 150mM NaCl, 4% SDS, 250U Benzonase) and the protein concentrations were determined using Bio-Rad DC™ Protein Assay.

### Click chemistry

Parasite lysates at 1mg/mL concentration were treated with freshly premixed click chemistry reaction cocktail [100µM capture reagent (TAMRA-Alkyne or Biotin-Alkyne; 10mM stock solution in DMSO), 1mM CuSO_4_ solution (50mM stock solution in MilliQ water), 1mM TCEP solution (50mM stock solution in MilliQ water) and 100µM TBTA (10mM stock solution in DMSO)] for 3 hours at room temperature. Proteins were precipitated by adding methanol (4 volumes), chloroform (1.5 volumes) and water (3 volumes) and collected by centrifugation at 16,000 g for 5 minutes. The protein precipitates were washed twice with methanol (10 volumes; centrifugation at 16,000 g for 5 minutes to collect the pellets) and the supernatants were discarded. The protein pellets were air-dried at room temperature for 20 minutes and stored in −80 °C freezer.

### In-gel fluorescence scanning

The air-dried protein pellets were suspended in resuspension buffer (4% SDS, 50mM HEPES pH 7.4, 150mM NaCl) to 1.33mg/mL final concentration. 4X Laemmli Sample Buffer (reducing) was added so that the final protein concentration was 1mg/mL. The samples were then boiled at 95 °C for 8 minutes and allowed to cool to room temperature. The proteins were resolved by SDS-PAGE (12% SDS Tris-HCl gels; 20µg of protein was loaded per gel lane). The gels were scanned for fluorescence labelling using a GE typhoon 5400 gel imager.

### Affinity enrichment

The air-dried protein pellets obtained after click chemistry and protein precipitation were dissolved in phosphate buffered saline (PBS) with 2% SDS to 5mg/mL concentration by sonication. In a typical experiment, 300µg of the parasite lysate after click chemistry and protein precipitation was resuspended in 50µL of the 2% SDS buffer. The samples were the diluted 20-fold with PBS so that the final SDS amount was 0.1%. The samples were centrifuged at 10,000 g for 5 minutes to remove insoluble debris and the clear soluble portion was used for the affinity enrichment. Typically, 30µL of NeutrAvidin-Agarose beads, freshly washed three times with 0.1% SDS buffer (0.1% SDS in PBS), were added to each of the sample and the mixtures were rotated on an end-over-end rotating shaker for 2 hours at room temperature. The beads were then washed 3 times with 1% SDS in PBS, 3 times with 6M urea in PBS, 3 times with PBS and once with 25mM TEAB buffer. Each washing was performed with 20 volumes of the washing solutions with respect to the bead volume and centrifugation of the beads between washing steps were carried out at 2,000 g for 1 minute at room temperature.

### On-bead reduction, alkylation and tryptic digestion

The beads after affinity enrichment were resuspended in 150µl of 25mM TEAB buffer and treated with 5mM TCEP (100mM stock solution in water) for 30 minutes at 50 °C. The beads were washed once with 25mM TEAB buffer and resuspended in 150µl of 25mM TEAB buffer and treated with 10mM CAA (200mM stock solution in water) in dark for 20 minutes at room temperature. The beads were again washed with 25mM TEAB buffer and resuspended in 200µl of fresh 50mM TEAB buffer and treated with 5µg of sequencing grade modified trypsin at 37 °C for 16 hours. The samples were centrifuged at 5,000 g for 5 minutes at room temperature to collect the supernatant. The beads were washed twice with 50% (v/v) ACN containing 0.1% (v/v) FA (50 µL for each wash) and mixed with the previous supernatant. The collected tryptic peptides were acidified to pH 3 using FA and evaporated to dryness. The peptides were then redissolved in 0.1% (v/v) FA solution in water and subjected to desalting on Pierce™ C-18 Spin Columns (Thermo Scientific; CN: 89873) following manufacturer’s instructions. The peptides were evaporated to complete dryness under a vacuum.

### iTRAQ 4-plex labelling

The iTRAQ labelling of the dried and desalted tryptic peptides were carried out using the iTRAQ® Reagents Multiplex Kit following manufacturer’s instructions with minor modifications. Briefly, the peptides were resuspended in equal amounts (30µL) of dissolution buffer (0.5M TEAB buffer supplied with the iTRAQ kit). 70µL of ethanol was added to each iTRAQ 4-plex reagent vial pre-equilibrated to room-temperature. The contents of the iTRAQ reagents vials were then carefully and quickly transferred to the respective vials of peptide digests (iTRAQ® 114 to the DMSO control; iTRAQ® 115 to 1 hour starvation; iTRAQ® 116 to 2 hour starvation and iTRAQ® 117 to 3 hour starvation). The labelling reactions were performed for 1.5 hours at room-temperature following which, the reactions were quenched with 100mM Tris base (1M stock solution) for 15 minutes at room-temperature. The samples labelled with the four different iTRAQ channels were then pooled into a fresh vial, and concentrated on speed-vac. The peptides were reconstituted in water with 0.1% (v/v) FA and 2% (v/v) ACN and subjected to desalting on C-18 Sep-Pak Classic cartridges (Waters; WAT051910) following manufacturer’s instructions. The eluted peptides were concentrated under vacuum and subjected to a second round of cleaning up on HILIC TopTip™ (PolyLC; TT200HIL) solid-phase extraction tips following manufacturer’s instructions. The eluted peptides were concentrated under vacuum and reconstituted in aqueous 0.1% (v/v) FA.

### LC-MS/MS analysis

The iTRAQ labelled tryptic peptides were separated on an Eksigent nanoLC 425 operating with a nano-flow module using Waters nanoEase HSS C18 T3 column (75µm x 250mm). A Waters trap column (Acquity M-Class Symmetry C18 5µm, 180µm x 20mm) was used prior to the main separating nano column. 2.5µL of sample peptides were separated by mobile phase A (0.1% formic acid in water) and mobile phase B (0.1% formic acid in ACN) at a flow rate of 300nL/minute over 110 minutes. The gradient used was the following, 5% B to 8% B (0 to 2 minutes), 8% B to 30% B (2 to 60 minutes), 30% B to 40% B (60 to 70 minutes), 40% B to 85% B (70 to 72 minutes), at 85% (72 to 78 minutes), 85% B to 5% B (78 to 80 minutes), at 5% B (80 to 110 minutes). The MS analysis was performed on a TripleTOF 5600 system (Sciex) in high-resolution mode. The MS acquisition time was set from gradient time 0 to 90 minutes and the MS spectra were collected in the mass range of 400 to 1600 m/z with 250ms accumulation time per spectrum. Further fragmentation of each MS spectrum occurred with a maximum of 30 precursors per cycle and 33ms minimum accumulation time for each precursor across the range of 100 to 1600 m/z and dynamic exclusion for 12sec. The MS/MS spectra were acquired in high sensitivity mode and the collision energies were increased by checking the ‘adjust CE when using iTRAQ reagents’ box in the acquisition method.

### iTRAQ quantitative proteomics MS data processing and analysis

For protein identification and quantification, the wiff files from the Sciex TripleTOF 5600 system were imported into MaxQuant [22] (version 1.6.3.4) with integrated Andromeda database search engine [23]. The MS/MS spectra were queried against *L. mexicana* sequences from UniProt KB (8,524 sequences). Database search employed the following parameters: Reporter ion MS2 with multiplicity 4plex iTRAQ, trypsin digestion with maximum 2 missed cleavages, oxidation of methionine and acetylation of protein N-termini as variable modifications, carbamidomethylation of cysteine as fixed modification, maximum number of modifications per peptide set at 5, minimum peptide length of 6, and protein FDR 0.01. Appropriate correction factors for the individual iTRAQ channels for both peptide N-terminal labelling and lysine side-chain labelling as per the iTRAQ Reagent Multiplex Kit were also configured into the database search. The proteinGroups.txt file from the MaxQuant search output was processed using Perseus software [24] (version 1.6.2.3). Potential contaminants, reverse sequences and sequences only identified by site were filtered off. A One-sample t-test was performed on the two replicates and only proteins with p ≤ 0.05 were retained. Additionally, only proteins with at least 2 unique peptides identified were retained. For each identified protein, ratios of the AHA labelled Reporter Intensity Corrected vs. DMSO control sample from the corresponding replicate experiment was calculated yielding the fold change (FC). The FCs obtained for each protein were transformed into log2 scale, and volcano plots were generated between the calculated significance (-Log P-value) and the obtained FC in log2 scale for each protein across the three different duration of starvation.

### Gene Ontology analysis

The GO terms (Molecular Function, Biological Process, and Cellular Component) significantly enriched in the NSPs relative to the predicted *L. mexicana* proteome were derived using Trytripdb.org [25]. REVIGO software [26] (revigo.irb.hr) was employed to refine and visualise the enriched GO terms.

## Results

### AHA is metabolically incorporated into NSPs in *L. mexicana* promastigotes

Although the BONCAT approach has been extensively applied in mammalian cells, reports are relatively few in lower eukaryotes. Therefore, we first decided to test if AHA is metabolically incorporated into NSPs in *L. mexicana* promastigotes. As AHA is a methionine surrogate, successful application of the BONCAT technique often requires depleting of L-methionine from the intracellular methionine reserves, and this is accomplished by maintaining the cells in a methionine-free medium for a short duration. We treated *L. mexicana* promastigotes in methionine-free Schneider’s medium with 10% dialysed FBS for 30 minutes prior to incubation with AHA in the same medium for 1 hour. Treatment of protein synthesis inhibitor, cycloheximide (CHX), was used as a control. The parasite lysates were subjected to click reaction with a TAMRA-Alkyne reagent (S1 Figure) and the proteins after precipitation and re-solubilisation were resolved on SDS-PAGE and profiled via in-gel fluorescence scanning (Fig 1B). As shown in Fig 1C, intense fluorescence labelling of the NSP was observed even at the lower concentration of 100µM AHA treatment. Labelling saturation was observed at the higher concentration of 1mM AHA treatment. Importantly, even for the high concentration of 1mM AHA treatment, concurrent CHX treatment significantly diminished the protein labelling, indicating that the fluorescently labelled proteins are indeed newly synthesised.

### Metabolic incorporation of AHA into the NSP of *L. mexicana* promastigotes is sensitive to starvation

In order to test whether the AHA incorporation can be used for the labelling of NSPs during starvation in *L. mexicana* promastigotes, we incubated the parasites in DPBS for different time durations prior to the AHA treatment. Maintaining the parasites in DBPS without serum ensures severe starvation [27]. As shown in Fig 2, starvation-time-dependent decrease in the fluorescent labelling intensity was observed in the in-gel fluorescence scans, indicating that the AHA incorporation can be used for the labelling of the NSPs under starvation in this parasite.

**Fig 2.**
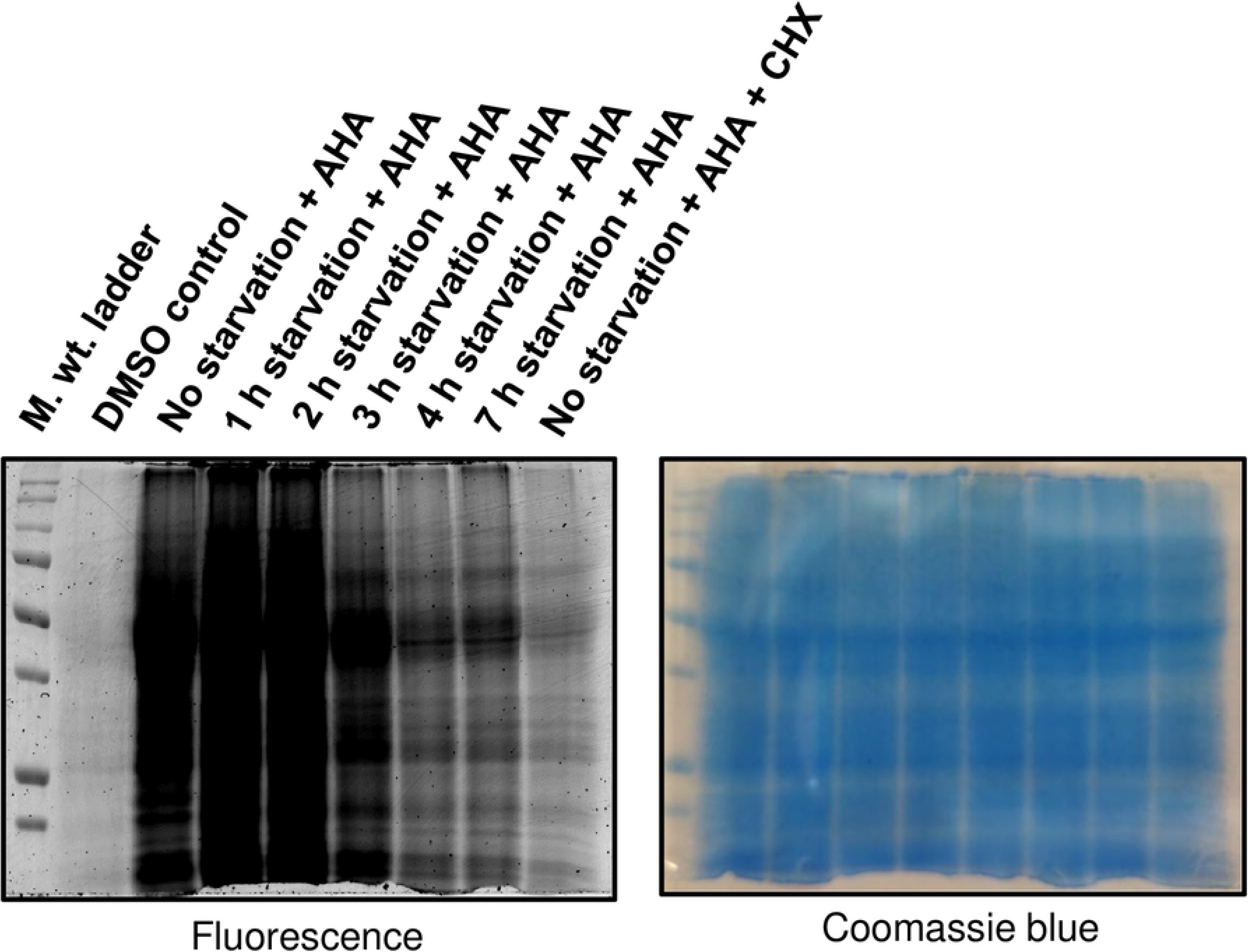
BONCAT in *L. mexicana* promastigotes under starvation. Promastigotes were cultured in methionine-free Schneider’s medium (30 minutes) prior to incubation in DPBS for different time periods (1 hour to 7 hours). The starved parasites were treated with AHA (50µM; lanes 3 to 9) or DMSO (control; lane 2) with (lane 9) or without CHX (10µM) for the last 1 hour of starvation and the NSPs were profiled by in-gel fluorescence scanning following click chemistry with a TAMRA-Alkyne. A Coomassie blue staining of the same gel that demonstrates even loading across the gel lanes is shown on the right panel.

### Development of a BONCAT-iTRAQ 4-plex workflow for quantitative proteomics MS profiling of the NSP during starvation in *L. mexicana* promastigotes

The in-gel fluorescence scanning only provides a qualitative information of the differential AHA labelling under starvation. In order to identify and generate a comparative quantitation of the NSPs at different time-points of starvation, we coupled the iTRAQ quantitative proteomics MS with the BONCAT. We used iTRAQ 4-plex labelling that enables comparison of 4 different experimental conditions in one experiment. As starvation beyond 3 hour duration was found to generate very little protein labelling in this parasite (Fig 2), we decided to compare the 1 hour, 2 hour and 3 hour time periods of starvation using quantitative proteomics. As shown in Fig 3, *L. mexicana* promastigotes, following the three different durations of incubation with DPBS, were treated with AHA to label the NSP. DMSO treatment instead of AHA was used as a control. The parasite lysates were subjected to click chemistry with a Biotin-Alkyne capture reagent (S1 Figure) and the labelled proteome were affinity enriched on NeutrAvidin-Agarose beads. The strong non-covalent interaction between biotin in the capture reagent and NeutrAvidin (K_d_ ≈ 10^−15^M) [28] permits stringent washing steps during the affinity enrichment protocol, enabling highly robust and selective pull-down of the labelled NSPs. After on-bead reduction, alkylation and tryptic digestion, the peptide digests were subjected to labelling with iTRAQ 4-plex reagents. The samples were then combined, desalted and analysed by nanoLC-MS/MS.

**Fig 3.**
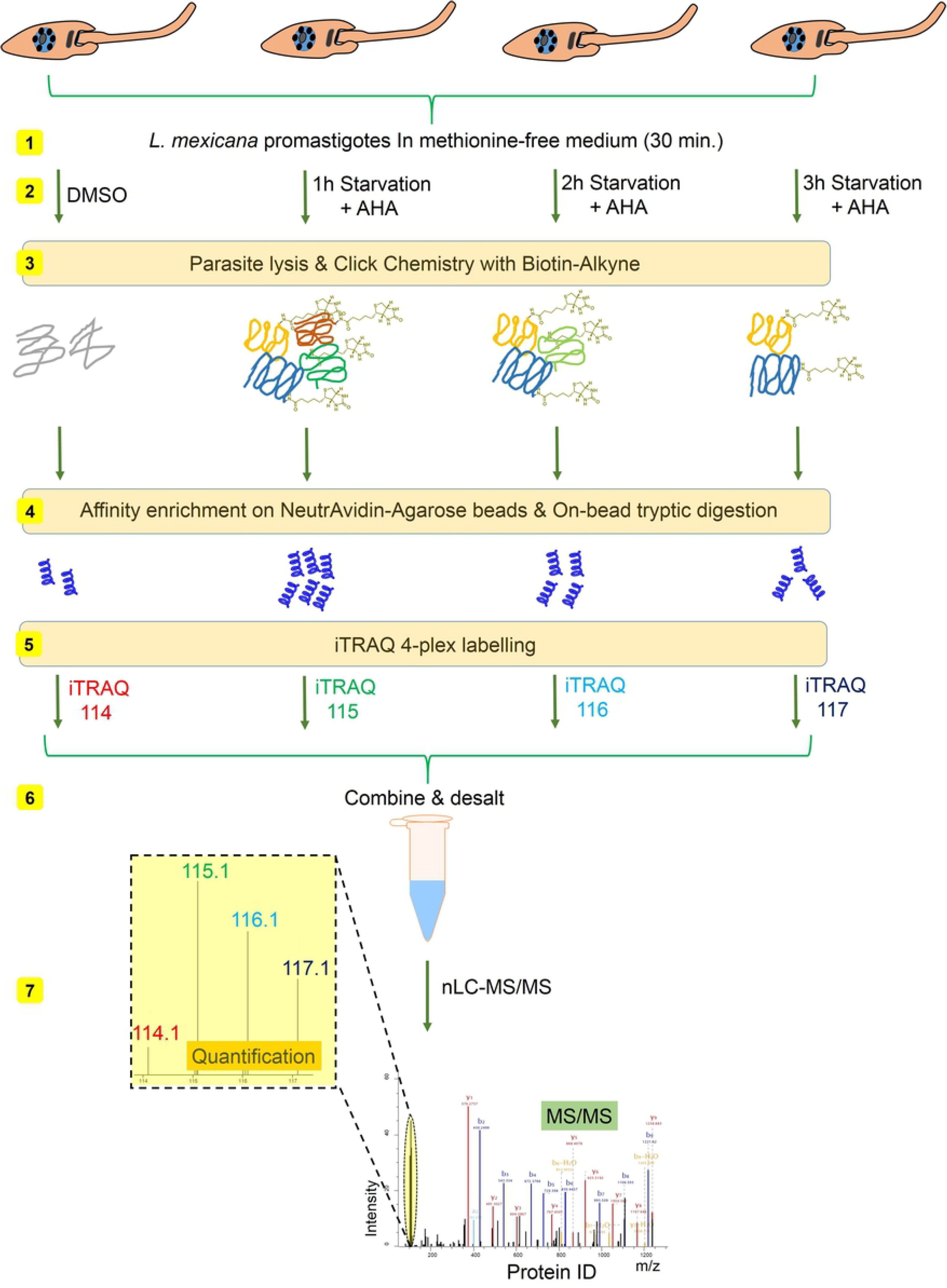
Schematic representation of the integrated BONCAT-iTRAQ 4-plex workflow used for profiling of NSPs of starving *L. mexicana* promastigotes. The NSPs in the parasites starved to 1 hour, 2 hour and 3 hour duration were tagged with AHA, following which the parasites were lysed, and the proteins were labelled using click reaction with Biotin-Alkyne. The labelled proteins were affinity enriched on NeutrAvidin beads, and following on-bead tryptic digestion, the released peptides were subjected to iTRAQ labelling. iTRAQ channel 114 was used for labelling the DMSO control sample, whilst channels 115, 116 and 117 were used respectively for labelling the NSPs at 1 hour, 2 hour and 3 hour starvation. The samples after pooling together were analysed by nanoLC-MS/MS.

### Identification and time-resolved quantitation of NSP in *L. mexicana* promastigotes during starvation

As shown in S1 Table, over 300 proteins were identified across the two replicate BONCAT-iTRAQ 4-plex experiments, of which 166 protein quantifications were statistically significant in a t-test analysis. For each of these NSPs, the iTRAQ reporter intensity ratio at each tested starvation duration to the DMSO control iTRAQ reporter intensity (iTRAQ 114 channel) within the same experiment was calculated. The observed fold change (FC) in abundance of each protein, after converting to log2 scale, was plotted against the significance in the t-test (-Log P-value). This enables filtering of the NSPs most significantly influenced by each tested duration of starvation (highlighted in blue in Fig 4A). A global decrease in the *de novo* protein synthesis was observed with increase in the duration of starvation. Functional annotation of the top-50 proteins by eggNOG database [29] revealed Translation, ribosomal structure and biogenesis and Posttranslational modification, protein turnover and chaperons along with Protein function unknown as the most abundant classifications (Fig 4B). The top-15 proteins with the highest changes in their abundance at each of the three tested duration of starvation are listed in the Table 1 (S2 Table, S3 Table and S4 Table report the top-50 NSPs identified at 1 hour, 2 hour and 3 hour starvation respectively). As shown in Table 1, many translation regulating proteins were observed among the top-ranking proteins at all three tested durations of starvation. Importantly, whilst the elongation initiation factor 2 alpha subunit, putative (Gene ID: LmxM.03.0980) and eukaryotic translation initiation factor 5A (Gene ID: LmxM.25.0720) were observed among the top-ranking proteins at the initial and intermediate stages of starvation (1 hour and 2 hours of starvation respectively), their relative abundance ranking among the NSPs went down at the later stage of starvation (3 hours duration). Thus, our data shows that at different durations of starvation in the parasite, a panel of important translation regulator proteins are expressed to different abundance levels.

**Table 1.**
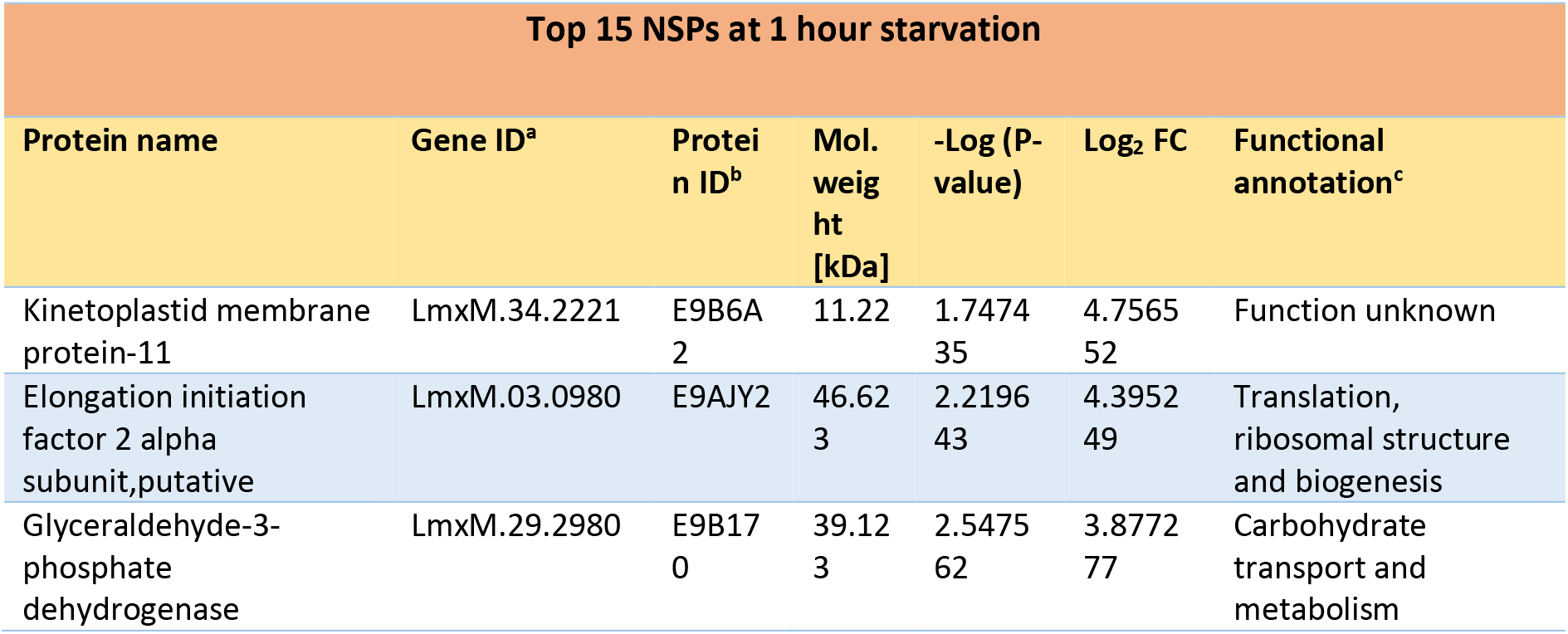
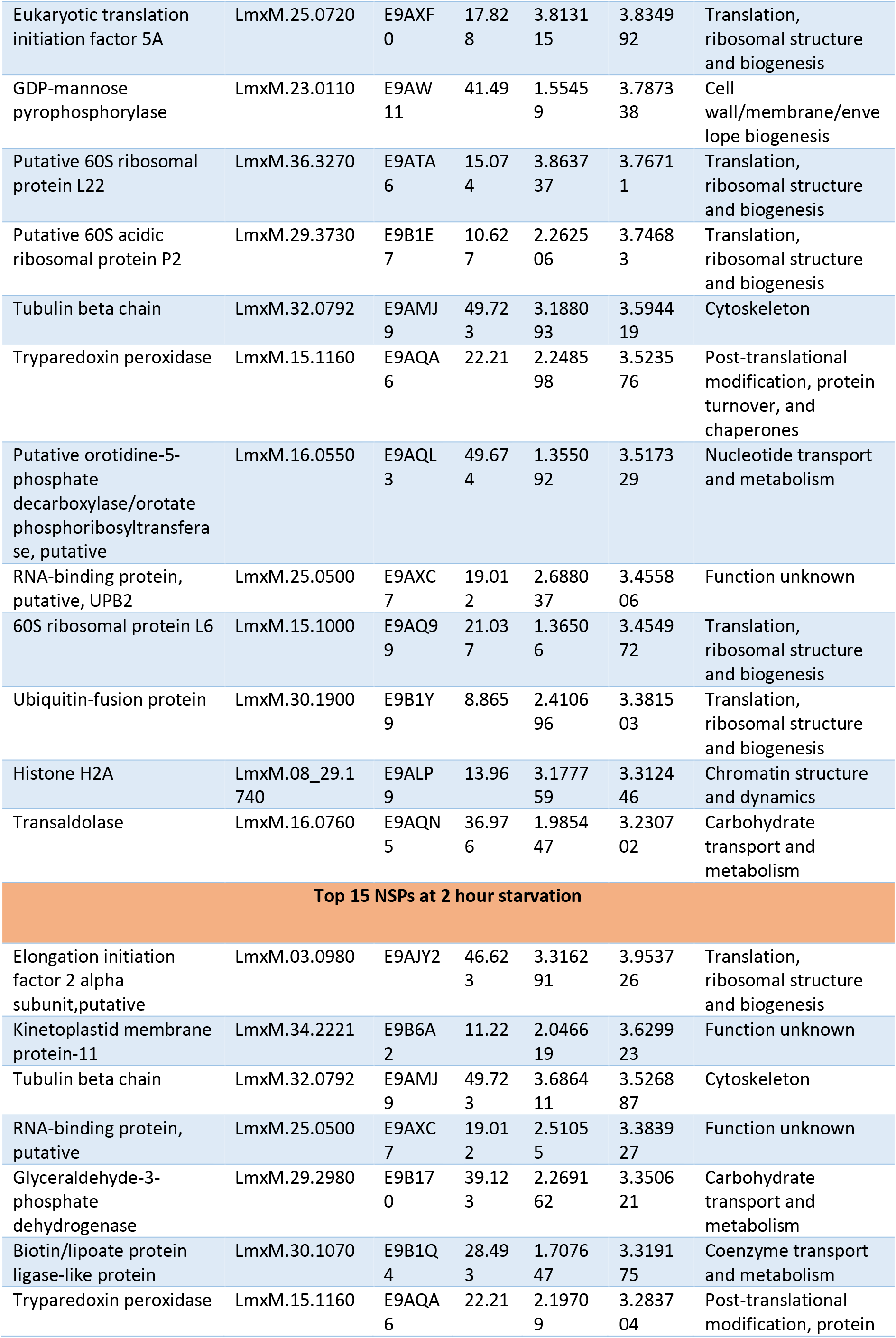
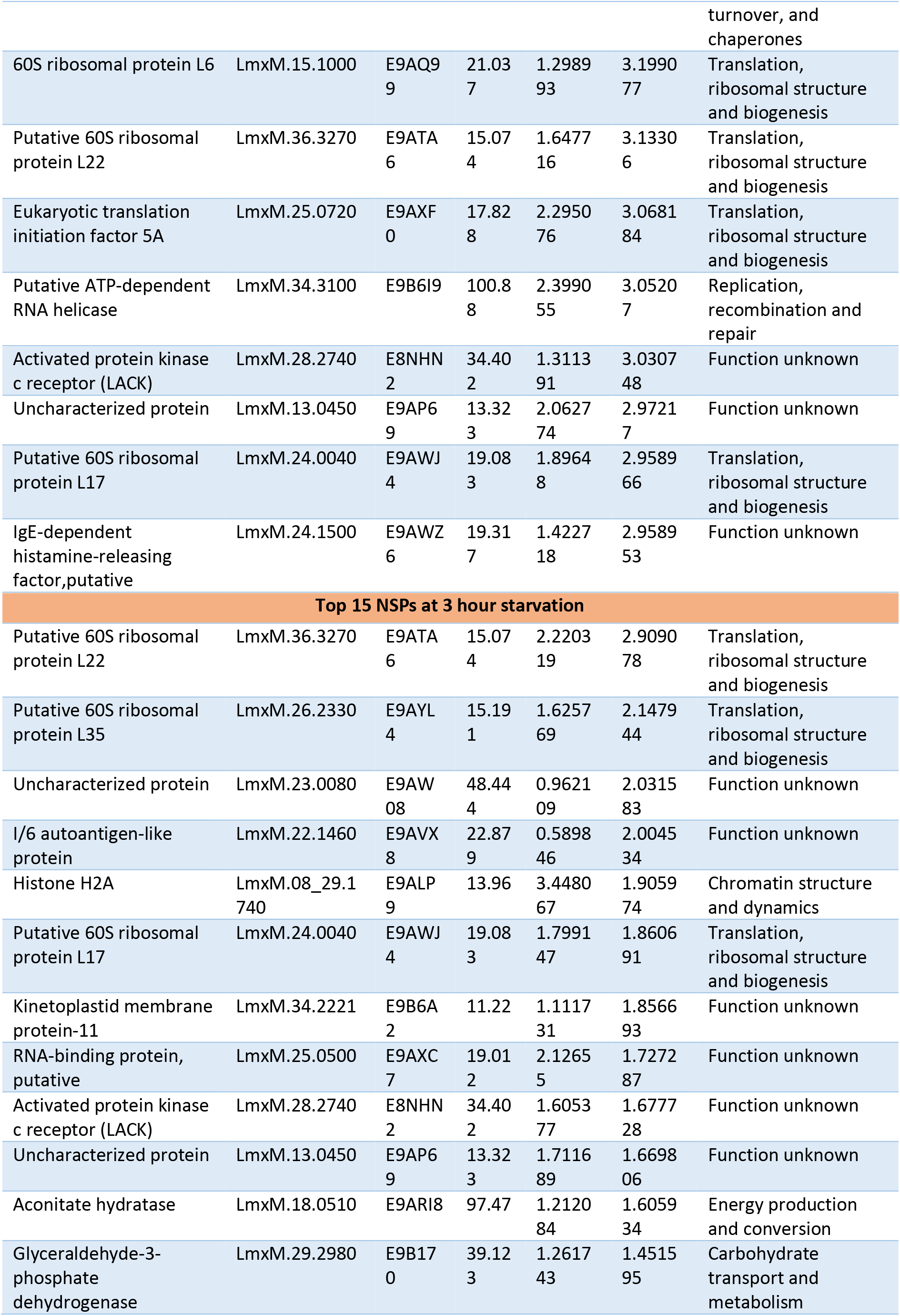
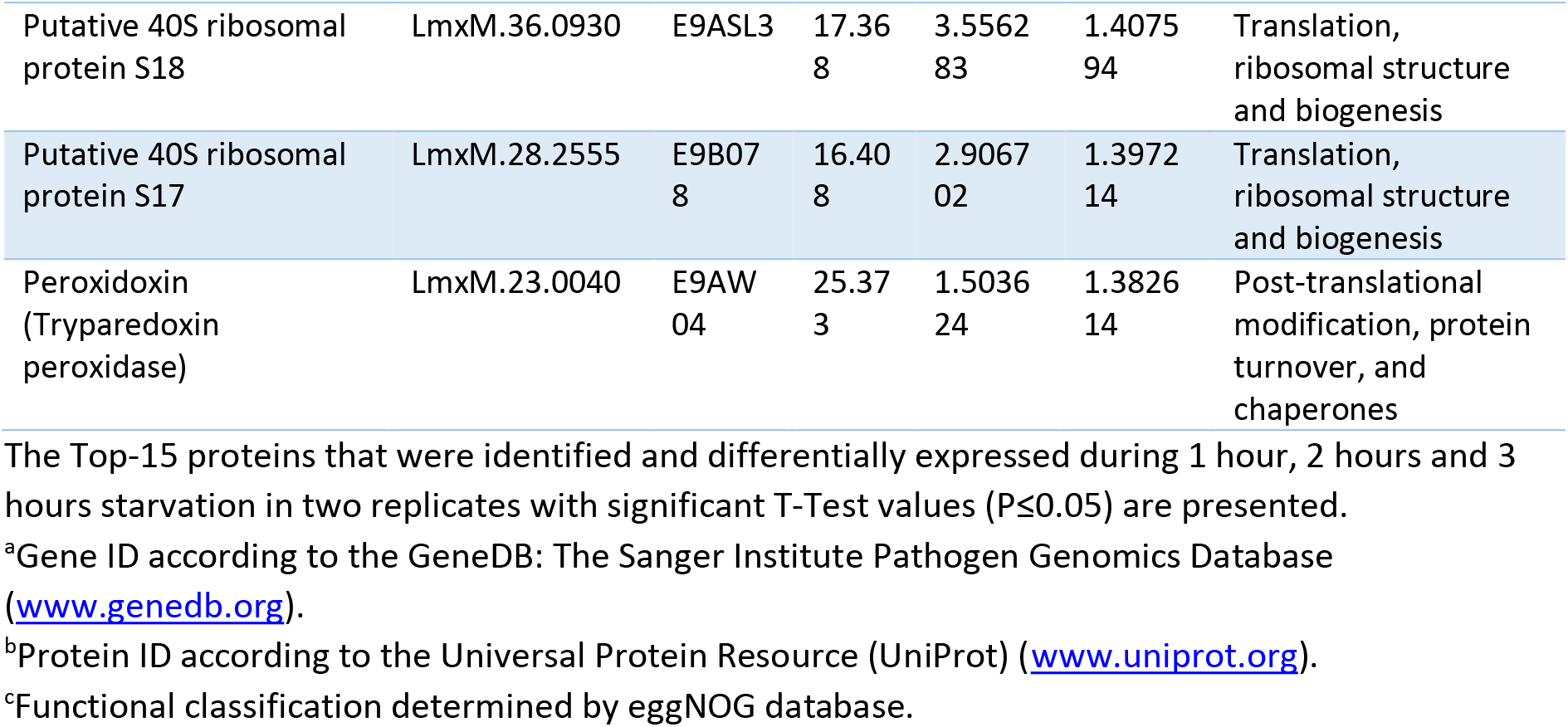
Top-15 NSPs identified in *L. mexicana* promastigotes upon starvation

**Fig 4.**
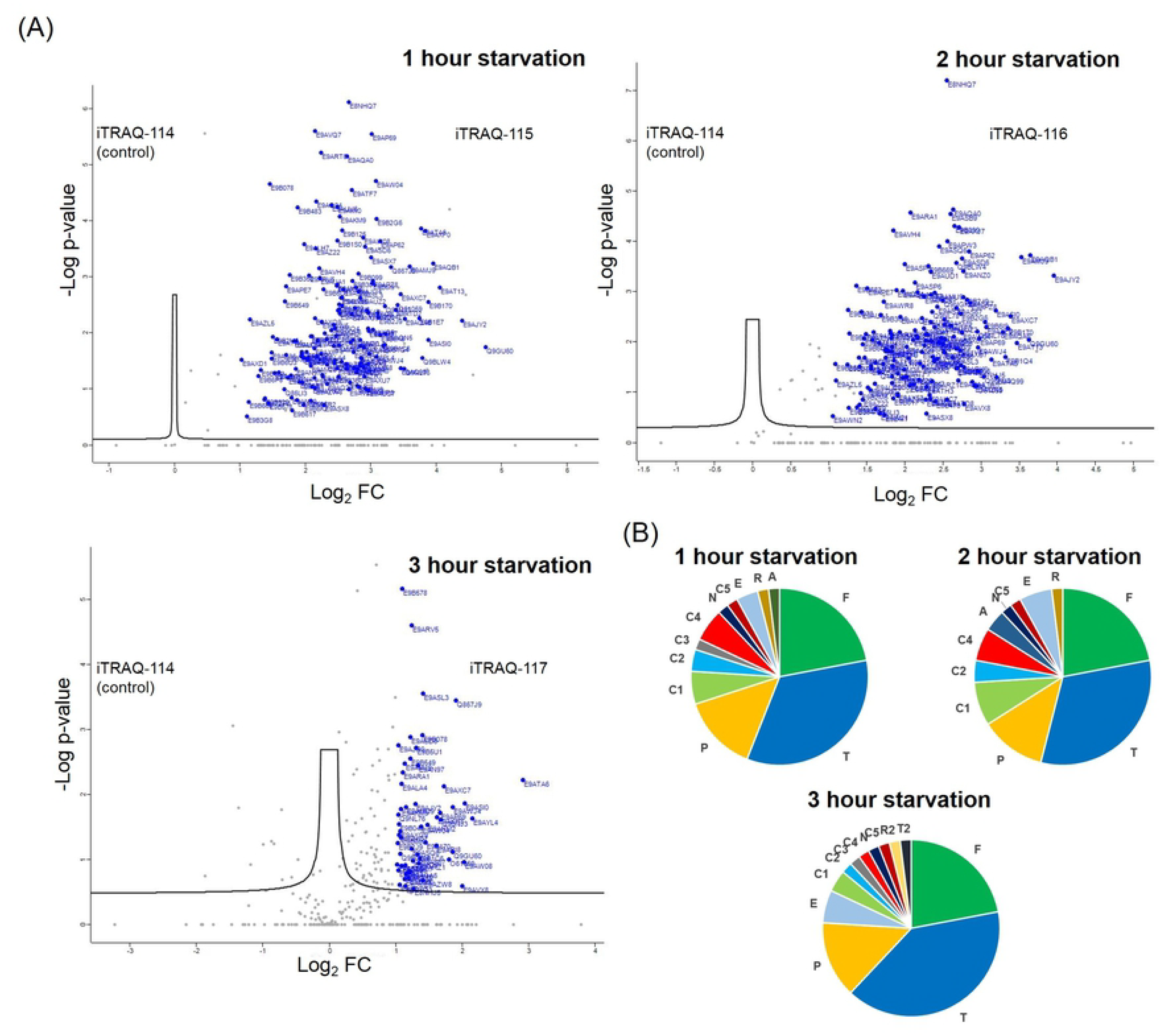
iTRAQ 4-plex quantitative proteomics MS profiling of NSPs of *L. mexicana* promastigotes during starvation. (**A**) Volcano plots of the NSP detected at the three durations of starvation. The significance of the iTRAQ reporter intensities obtained for each NSP at each tested duration of starvation across two replicates as −Log P-values was plotted against the observed fold change (FC) in abundance in log2 scale. Proteins with a log2 FC of more than 1 with significant iTRAQ quantifications are highlighted in blue. (**B**) Functional annotation pie chart of the top-50 NSPs. The letter codes used for the functional categories are the following. (T) Translation, ribosomal structure and biogenesis; (F) Function unknown; (P) Post-translational modification, protein turnover, and chaperones; (A) Amino acid transport and metabolism; (C1) Carbohydrate transport and metabolism; (C2) Coenzyme transport and metabolism; (C3) Chromatin structure and dynamics; (C4) Cytoskeleton; (C5) Cell wall/membrane/envelope biogenesis; (R) Replication, recombination and repair; (R2) RNA processing and modification; (N) Nucleotide transport and metabolism; (T2) Transcription; (E) Energy production and conversion.

### Gene Ontology (GO) analysis of the NSPs in *L. mexicana* promastigotes during starvation

Biological Process GO term enrichment analysis (Fig 5A) of the complete 166 statistically significant protein IDs of the NSPs revealed translation (P value 7.82e^−68^; 86 entries) and peptide biosynthetic process (P value 2.01e^−67^; 86 entries) as the most significantly enriched terms. Gene expression (P value 1.04e^−50^; 91 entries) was also among highly enriched terms. Ribosome (P value 4.47e^−68^; 76 entries) and ribonucleoprotein complex (P value 6.66e^−61^; 76 entries) were the most significantly enriched Cellular Component GO terms (Fig 5B). Similarly, Molecular Function GO term analysis (Fig 5C) revealed structural constituent of ribosome (P value 1.22e^−68^; 76 entries), structural molecular activity (P value 2.11e^−65^; 76 entries), RNA binding (P value 3.66e^−10^; 28 entries) and unfolded protein binding (P value 5.22e^−9^; 12 entries) as the most significantly enriched terms. The GO analyses clearly indicate the high specificity of the identified NSP towards regulation of protein synthesis in the ribosome relative to the available data of whole cell proteome of the parasite (Tritrypdb.org).

**Fig 5.**
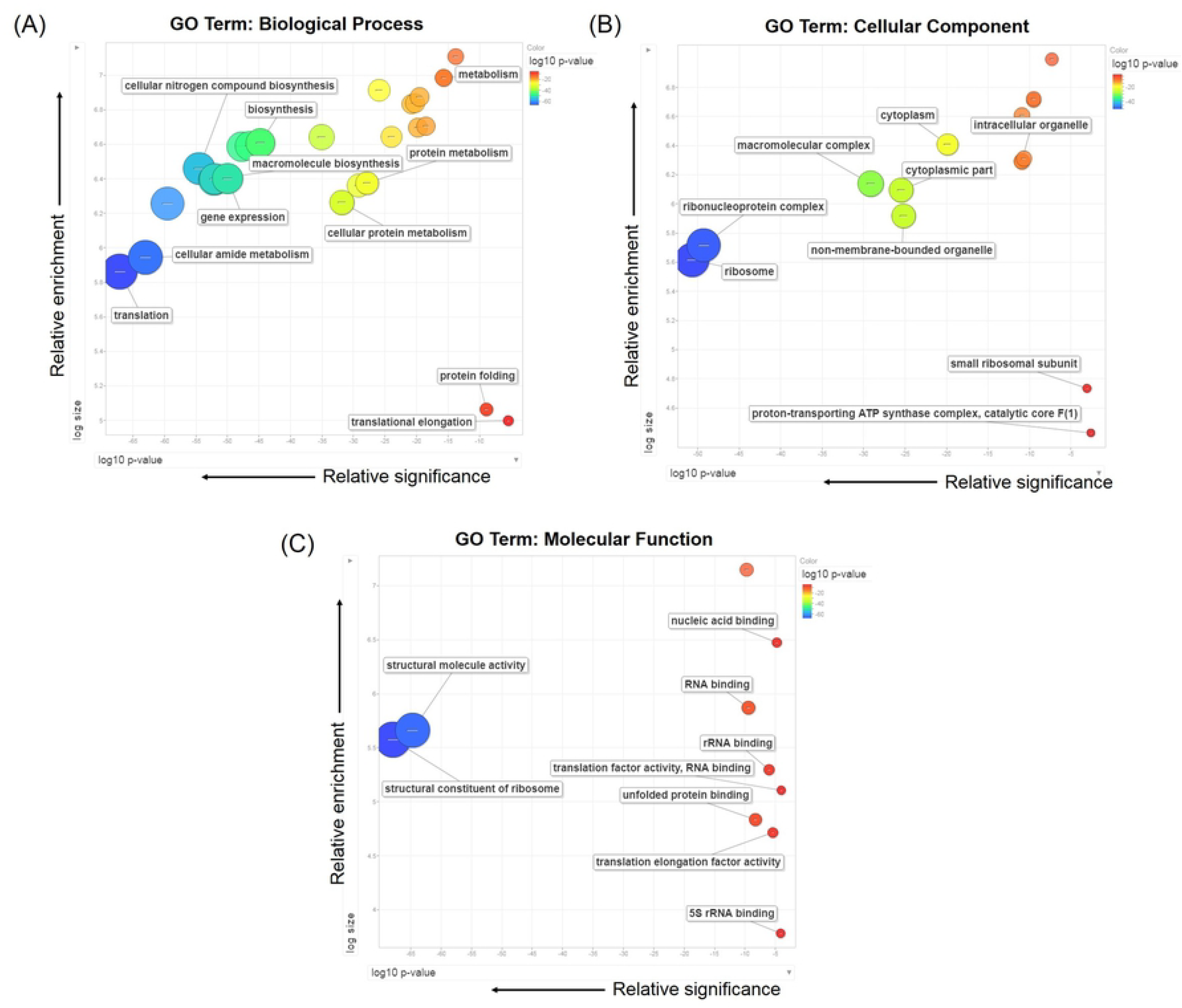
Gene Ontology Term enrichment of the 166 NSPs relative to the predicted *L. mexicana* whole proteome. (**A**) GO term enrichment for Biological Process (**B**) GO term enrichment for Cellular Component, and (**C**) GO term enrichment for Molecular Function. The GO terms were refined and visualised using REVIGO software.

## Discussion

Quantitative proteomics profiling of NSPs during severe starvation in the *Leishmania* parasites require a methodology that is robust and sensitive to distinguish the lower-abundance NSPs from the pre-existing pool of the parasite proteome. We reasoned that a workflow that combines the BONCAT technique with a peptide-level labelling-based quantitative proteomics MS technique such as iTRAQ or TMT labelling could be developed to meet this requirement. The high efficiency and bio-orthogonality of the click reaction could ensure robust and preferential enrichment of the NSPs. Besides, AHA treatment has been proven to be non-toxic as it does not cause significant protein misfolding or alterations in the global protein turn over [8,30]. Similarly, the iTRAQ quantitative proteomics MS offers a powerful technology for comparative proteomics. It enables highly sensitive and reliable quantitation of protein abundance changes across multiple experimental conditions. iTRAQ, and other similar labelling-based quantitative MS techniques offer far more reliable and reproducible proteome quantitation than the different versions of spectral counting or precursor ion signal intensity-based label-free quantitative MS [31,32]. The sample multiplexing of iTRAQ method provides an additional benefit of peptide signal pooling effect, which increases the sensitivity of detection; a particularly useful feature in the starvation experiment, a context where the global protein synthesis is significantly lowered. Thus, the unique combination of BONCAT approach and iTRAQ quantitative proteomics MS provided a workflow that is robust and sensitive to profiling the NSPs of *L. mexicana* promastigotes under severe starvation.

Although the alternative, ribosome profiling [33,34] is emerging as a powerful method for global profiling of protein translation, MS-based proteomics, comparatively, provides a more direct, and therefore more reliable, readout of the cellular proteome and its changes under different perturbations [35]. Proteins are more robust during sample handling, whilst every step in the experimental protocols of ribosome profiling from cell lysis to nuclease digestion to library generation is likely to cause distortions in the data [36]. The use of translational inhibitors during ribosome profiling is also known to affect the local distribution of ribosomes on mRNAs [37]. Additionally, false readout of translation due to contaminating ribosomal RNA (rRNA) fragments is a common occurrence in ribosome profiling [38]. In a starvation condition, when the global translation levels are low, the rRNA contaminating fragments could significantly compromise the ribosome footprint sequencing space [33]. Our BONCAT-iTRAQ MS approach in *Leishmania* provides a powerful alternative to the ribosome profiling in the protozoa, and the method in *L. mexicana* promastigotes enabled direct profiling of the NSPs and their relative quantitative changes in abundance under starvation in a time-dependent manner.

In higher eukaryotes, the eukaryotic initiation factor 2 alpha (eIF2α) serves as an essential component for protein synthesis. It also acts as a key translation regulator during stress conditions including nutrient starvation [39]. The eIF2α-mediated translational regulation has been reported to facilitate differentiation in *Leishmania* parasites [40]. The observation of the elongation initiation factor 2 alpha subunit, putative (Gene ID: LmxM.03.0980) along with several other translation-facilitating proteins among the top-ranking proteins in the early and intermediate stages of starvation in *L. mexicana* promastigotes compared to the observed lower ranking of these proteins in the later stage of starvation indicates a starvation time-dependent differential regulation of protein synthesis in the parasite. Some of the top-ranking proteins identified in this study are known to be important from a disease-tackling point of view. For instance, the kinetoplast membrane protein-11, a protein that is conserved in all kinetoplastids, has been characterised as a virulence factor in *L. amazonensis* infection and is investigated as a vaccine candidate [41]. Importantly, the expression of this protein has been previously reported to be upregulated during metacyclogenesis [42]. Another important protein, activated protein kinase c receptor (LACK), has been reported to act as a T-cell epitope, and was proposed as yet another potential candidate for vaccine development [43]. Another top-ranking protein, tryparedoxin peroxidase, is an important enzyme the parasite relies on for detoxifying reactive oxygen species [44]. This protein has been found to be upregulated in amphotericin B-resistant isolates [45] and antimony-resistant isolates of *Leishmania supp*. [46], indicating its possible role in drug resistance. Yet another top-ranking protein glyceraldehyde-3-phosphate dehydrogenase has been reported to be required for survival of *L. donovani* in visceral organs [47].

Our results indicate that Translation, ribosome structure and biogenesis and Posttranslational modifications, protein turnover and chaperons were among the most representative enriched functional annotations of the NSPs identified. This is in congruence with the previous finding that *Leishmania* exerts an increased level of control on translation during stress conditions [40]. A higher level control on translation is expected under starvation as translation is energetically a costly process for the cell [48], and therefore the parasite has to rely on an increased level of control on translation, and potentially posttranslational mechanisms as well, for conserving the available limited nutrient resources, and to optimise and appropriately regulate protein synthesis to avoid generating toxic protein forms. This is the first study that comprehensively and quantitatively profiled the NSPs during starvation in *Leishmania*. It is, however, important to note that despite the recent advancements in the genome sequencing of several *Leishmania* strains, a major portion of the predicted parasite proteome remain functionally unannotated and termed uncharacterised. Nevertheless, bioinformatics methods such as protein-protein interaction mapping [49], domain identification [50] and structural homology modelling [51] are making advancements in the protein functional annotation efforts. Therefore, we believe that along with future developments in more detailed functional characterisation of the *Leishmania* proteome, our results will provide additional insights into the molecular mechanisms involved in regulating the gene expression under severe starvation in the protozoan. Regulation of protein synthesis in kinetoplastids is currently poorly understood. Our method introduces a powerful platform for studying the protein synthesis in the parasites in a temporally resolved, quantitative and high-throughput manner. It is anticipated that our methodology will find wide-spread applications in the kinetoplastida parasites and in the broader area of NTD, and the results from this study will serve as a starting point for future studies to unravel the starvation-adaptation mechanisms in different life cycle stages in these parasites.

## Acknowledgements

We acknowledge stimulating discussions with Professor Patrick G. Steel, Department of Chemistry, Durham University, UK and Associate Professor Steven Cobb, Department of Chemistry, Durham University, UK. Special thanks to Dr Adrian Brown, Proteomics Facility, Department of Biosciences, Durham University, UK for technical support on HILIC solid-phase extraction and for the nanoLC-MS/MS runs.

## Author Contributions

Conceived of the study and designed experiments: KK. Performed experiments: KK. Oversaw project management: PWD, KK. Analysed the data: KK. Contributed reagents and materials: PWD, KK. Wrote the paper: KK. Reviewed the paper: PWD.

## Supporting Information

**S1 Fig. Chemical structure of the capture reagents used.** (**A**) 5-TAMRA-Alkyne used for click chemistry followed by in-gel fluorescence imaging. (**B**) Acetylene-PEG4-Biotin used for click chemistry followed by affinity enrichment and iTRAQ proteomics MS. (TIFF)

**S2 Fig. Gene Ontology word cloud of the NSPs identified in *L. mexicana* promastigotes during starvation.** (**A**) Biological Process GO term enrichment word cloud (**B**) Cellular Component GO term enrichment word cloud, and (**C**) Molecular Function GO term enrichment word cloud. (TIFF)

**S1 Table. LC-MS/MS protein identification and quantification output.** The complete list of proteins identified in two replicate iTRAQ 4-plex labelling experiments along with the corrected reported intensities of the four iTRAQ channels for each protein in the two experiments following MaxQuant-Perseus database search and data processing. The corrected reporter intensities of each iTRAQ channel is presented as a fold change (FC) in log2 scale from the DMSO control 114 channel. The symbol NaN indicates a non-valid value resulting from missing reporter ion signals. T-test significant (p ≤ 0.05) entries are indicated with a + sign and only those proteins that are both significant and with 2 or more identified unique peptides were used for subsequent bioinformatic analysis. (XLSX)

**S2 Table. Top-50 NSPs identified at 1 hour starvation.** The proteins are listed in the descending order of their observed FC in abundance values in log2 scale. (PDF)

**S3 Table. Top-50 NSPs identified at 2 hour starvation.** The proteins are listed in the descending order of their observed FC in abundance values in log2 scale. (PDF)

**S4 Table. Top-50 NSPs identified at 3 hour starvation.** The proteins are listed in the descending order of their observed FC in abundance values in log2 scale. (PDF)

## References

1. Alvar J, Velez ID, Bern C, Herrero M, Desjeux P, Cano J, et al. Leishmaniasis worldwide and global estimates of its incidence. PLoS One. 2012;7(5):e35671. PMID: 22693548.

2. De Pablos LM, Ferreira TR, Walrad PB. Developmental differentiation in Leishmania lifecycle progression: post-transcriptional control conducts the orchestra. Curr Opin Microbiol. 2016;34:82–9. PMID: 27565628.

3. Carter NS, Yates PA, Gessford SK, Galagan SR, Landfear SM, Ullman B. Adaptive responses to purine starvation in Leishmania donovani. Mol Microbiol. 2010;78(1):92–107. PMID: 20923417.

4. Spath GF, Drini S, Rachidi N. A touch of Zen: post-translational regulation of the Leishmania stress response. Cell Microbiol. 2015;17(5):632–8. PMID: 25801803.

5. Martin JL, Yates PA, Soysa R, Alfaro JF, Yang F, Burnum-Johnson KE, et al. Metabolic reprogramming during purine stress in the protozoan pathogen Leishmania donovani. PLoS Pathog. 2014;10(2):e1003938. PMID: 24586154.

6. Serafim TD, Figueiredo AB, Costa PAC, Marques-Da-Silva EA, Goncalves R, de Moura SAL, et al. Leishmania Metacyclogenesis Is Promoted in the Absence of Purines. PLoS Negl Trop Dis. 2012;6(9):e1833. PMID: 23050028.

7. Louradour I, Monteiro CC, Inbar E, Ghosh K, Merkhofer R, Lawyer P, et al. The midgut microbiota plays an essential role in sand fly vector competence for Leishmania major. Cell Microbiol. 2017;19(10):e12755. PMID: 28580630.

8. Dieterich DC, Link AJ, Graumann J, Tirrell DA, Schuman EM. Selective identification of newly synthesized proteins in mammalian cells using bioorthogonal noncanonical amino acid tagging (BONCAT). Proc Natl Acad Sci U S A. 2006;103(25):9482–7. PMID: 16769897.

9. Dieterich DC, Lee JJ, Link AJ, Graumann J, Tirrell DA, Schuman EM. Labeling, detection and identification of newly synthesized proteomes with bioorthogonal non-canonical amino-acid tagging. Nat Protoc. 2007;2(3):532–40. PMID: 17406607.

10. Ross PL, Huang YLN, Marchese JN, Williamson B, Parker K, Hattan S, et al. Multiplexed protein quantitation in Saccharomyces cerevisiae using amine-reactive isobaric tagging reagents. Mol Cell Proteomics. 2004;3(12):1154–69. PMID: 15385600.

11. Wiese S, Reidegeld KA, Meyer HE, Warscheid B. Protein labeling by iTRAQ: A new tool for quantitative mass spectrometry in proteome research. Proteomics. 2007;7(3):340–50. PMID: 17177251.

12. Clayton C, Shapira M. Post-transcriptional regulation of gene expression in trypanosomes and leishmanias. Mol Biochem Parasitol. 2007;156(2):93–101. PMID: 17765983.

13. Haile S, Papadopoulou B. Developmental regulation of gene expression in trypanosomatid parasitic protozoa. Curr Opin Microbiol. 2007;10(6):569–77. PMID: 18177626.

14. Kramer S. Developmental regulation of gene expression in the absence of transcriptional control: The case of kinetoplastids. Mol Biochem Parasitol. 2012;181(2):61–72. PMID: 22019385.

15. Lahav T, Sivam D, Volpin H, Ronen M, Tsigankov P, Green A, et al. Multiple levels of gene regulation mediate differentiation of the intracellular pathogen Leishmania. FASEB J. 2011;25(2):515–25. PMID: 20952481.

16. de Pablos LM, Ferreira TR, Dowle AA, Forrester S, Parry E, Newling K, et al. The mRNA-bound Proteome of Leishmania mexicana: Novel Genetic Insight into an Ancient Parasite. Mol Cell Proteomics. 2019;18(7):1271–84. PMID: 30948621.

17. Walther TC, Mann M. Mass spectrometry-based proteomics in cell biology. J Cell Biol. 2010;190(4):491–500. PMID: 20733050.

18. Presolski SI, Hong VP, Finn MG. Copper-Catalyzed Azide-Alkyne Click Chemistry for Bioconjugation. Curr Protoc Chem Biol. 2011;3(4):153–62. PMID: 22844652.

19. Thompson A, Schafer J, Kuhn K, Kienle S, Schwarz J, Schmidt G, et al. Tandem mass tags: a novel quantification strategy for comparative analysis of complex protein mixtures by MS/MS. Anal Chem. 2003;75(8):1895–904. PMID: 12713048.

20. Mertins P, Udeshi ND, Clauser KR, Mani DR, Patel J, Ong SE, et al. iTRAQ Labeling is Superior to mTRAQ for Quantitative Global Proteomics and Phosphoproteomics. Mol Cell Proteomics. 2012;11(6):M111.014423. PMID: 22210691.

21. Kalesh K, Lukauskas S, Borg AJ, Snijders AP, Ayyappan V, Leung AKL, et al. An Integrated Chemical Proteomics Approach for Quantitative Profiling of Intracellular ADP-Ribosylation. Sci Rep. 2019;9(1):6655. PMID: 31040352.

22. Cox J, Mann M. MaxQuant enables high peptide identification rates, individualized p.p.b.-range mass accuracies and proteome-wide protein quantification. Nat Biotechnol. 2008;26(12):1367–72. PMID: 19029910.

23. Cox J, Neuhauser N, Michalski A, Scheltema RA, Olsen JV, Mann M. Andromeda: A Peptide Search Engine Integrated into the MaxQuant Environment. J Proteome Res. 2011;10(4):1794–805. PMID: 21254760.

24. Tyanova S, Temu T, Sinitcyn P, Carlson A, Hein MY, Geiger T, et al. The Perseus computational platform for comprehensive analysis of (prote)omics data. Nat Methods. 2016;13(9):731–40. PMID: 27348712.

25. Aslett M, Aurrecoechea C, Berriman M, Brestelli J, Brunk BP, Carrington M, et al. TriTrypDB: a functional genomic resource for the Trypanosomatidae. Nucleic Acids Res. 2010;38:D457–62. PMID: 19843604.

26. Supek F, Bosnjak M, Skunca N, Smuc T. REVIGO Summarizes and Visualizes Long Lists of Gene Ontology Terms. PloS One. 2011;6(7):e21800. PMID: 21789182.

27. Besteiro S, Williams RA, Morrison LS, Coombs GH, Mottram JC. Endosome sorting and autophagy are essential for differentiation and virulence of Leishmania major. J Biol Chem. 2006;281(16):11384–96. PMID: 16497676.

28. Green NM. Avidin and Streptavidin. Method Enzymol. 1990;184:51–67. PMID: 2388586.

29. Powell S, Forslund K, Szklarczyk D, Trachana K, Roth A, Huerta-Cepas J, et al. eggNOG v4.0: nested orthology inference across 3686 organisms. Nucleic Acids Res. 2014;42(D1):D231–9. PMID: 24297252.

30. Hinz FI, Dieterich DC, Schuman EM. Teaching old NCATs new tricks: using non-canonical amino acid tagging to study neuronal plasticity. Curr Opin Chem Biol. 2013;17(5):738–46. PMID: 23938204.

31. Lai X, Wang L, Witzmann FA. Issues and applications in label-free quantitative mass spectrometry. Int J Proteomics. 2013;2013:756039. PMID: 23401775.

32. Rauniyar N, Yates JR, 3rd. Isobaric labeling-based relative quantification in shotgun proteomics. J Proteome Res. 2014;13(12):5293–309. PMID: 25337643.

33. Ingolia NT, Ghaemmaghami S, Newman JRS, Weissman JS. Genome-Wide Analysis in Vivo of Translation with Nucleotide Resolution Using Ribosome Profiling. Science. 2009;324(5924):218–23. PMID: 19213877.

34. Bifeld E, Lorenzen S, Bartsch K, Vasquez JJ, Siegel TN, Clos J. Ribosome Profiling Reveals HSP90 Inhibitor Effects on Stage-Specific Protein Synthesis in Leishmania donovani. mSystems. 2018;3(6):e00214–18. PMID: 30505948.

35. Liu TY, Huang HH, Wheeler D, Xu Y, Wells JA, Song YS, et al. Time-Resolved Proteomics Extends Ribosome Profiling-Based Measurements of Protein Synthesis Dynamics. Cell Syst. 2017;4(6):636–44 e9. PMID: 28578850.

36. Brar GA, Weissman JS. Ribosome profiling reveals the what, when, where and how of protein synthesis. Nat Rev Mol Cell Biol. 2015;16(11):651–64. PMID: 26465719.

37. Gerashchenko MV, Gladyshev VN. Translation inhibitors cause abnormalities in ribosome profiling experiments. Nucleic Acids Res. 2014;42(17):e134. PMID: 25056308.

38. Gerashchenko MV, Lobanov AV, Gladyshev VN. Genome-wide ribosome profiling reveals complex translational regulation in response to oxidative stress. Proc Natl Acad Sci U S A. 2012;109(43):17394–9. PMID: 23045643.

39. Baird TD, Wek RC. Eukaryotic Initiation Factor 2 Phosphorylation and Translational Control in Metabolism. Adv Nutr. 2012;3(3):307–21. PMID: 22585904.

40. Cloutier S, Laverdiere M, Chou MN, Boilard N, Chow C, Papadopoulou B. Translational Control through eIF2alpha Phosphorylation during the Leishmania Differentiation Process. PloS One. 2012;7(5):e35085. PMID: 22693545.

41. de Mendonca SCF, Cysne-Finkelstein L, Matos DCD. Kinetoplastid membrane protein-11 as a vaccine candidate and a virulence factor in Leishmania. Front Immunol. 2015;6:524. PMID: 26528287.

42. Matos DC, Faccioli LA, Cysne-Finkelstein L, Luca PM, Corte-Real S, Armoa GR, et al. Kinetoplastid membrane protein-11 is present in promastigotes and amastigotes of Leishmania amazonensis and its surface expression increases during metacyclogenesis. Mem Inst Oswaldo Cruz. 2010;105(3):341–7. PMID: 20512252.

43. Sinha S, Kumar A, Sundaram S. Acomprehensive analysis of LACK (Leishmania homologue of receptors for activated C kinase) in the context of Visceral Leishmaniasis. Bioinformation. 2013;9(16):832–7. PMID: 24143055.

44. Iyer JP, Kaprakkaden A, Choudhary ML, Shaha C. Crucial role of cytosolic tryparedoxin peroxidase in Leishmania donovani survival, drug response and virulence. Mol Microbiol. 2008;68(2):372–91. PMID: 18312262.

45. Suman SS, Equbal A, Zaidi A, Ansari MY, Singh KP, Singh K, et al. Up-regulation of cytosolic tryparedoxin in Amp B resistant isolates of Leishmania donovani and its interaction with cytosolic tryparedoxin peroxidase. Biochimie. 2016;121:312–25. PMID: 26743980.

46. Andrade JM, Murta SM. Functional analysis of cytosolic tryparedoxin peroxidase in antimony-resistant and -susceptible Leishmania braziliensis and Leishmania infantum lines. Parasit Vectors. 2014;7:406. PMID: 25174795.

47. Zhang WW, McCall LI, Matlashewski G. Role of cytosolic glyceraldehyde-3-phosphate dehydrogenase in visceral organ infection by Leishmania donovani. Eukaryot Cell. 2013;12(1):70–7. PMID: 23125352.

48. Lynch M, Marinov GK. The bioenergetic costs of a gene. Proc Natl Acad Sci U S A. 2015;112(51):15690–5. PMID: 26575626.

49. Legrain P, Wojcik J, Gauthier JM. Protein--protein interaction maps: a lead towards cellular functions. Trends Genet. 2001;17(6):346–52. PMID: 11377797.

50. Feldman HJ. Identifying structural domains of proteins using clustering. BMC Bioinformatics. 2012;13:286. PMID: 23116496.

51. Waterhouse A, Bertoni M, Bienert S, Studer G, Tauriello G, Gumienny R, et al. SWISS-MODEL: homology modelling of protein structures and complexes. Nucleic Acids Res. 2018;46(W1):W296–303. PMID: 29788355.

